# Midgut aminopeptidase N expression profile in Castor semilooper during sublethal Cry toxin exposure

**DOI:** 10.1101/849612

**Authors:** Vinod K. Chauhan, Narender K. Dhania, Vadthya Lokya, Bhoopal Bhuvanachandra, Kollipara Padmasree, Aparna Dutta-Gupta

**Author notes:** Corresponding author, /.

## Abstract

Midgut of lepidopteran larvae is a multifunctional tissue, which performs roles in digestion, absorption, immunity; transmission of pathogens and interaction with ingested various molecules. The proteins localized at the inner apical brush border membrane are primarily digestive proteases but some of them like aminopeptidase N, alkaline phosphatase, cadherins, ABC transporter C2 etc. interact with Crystal (Cry) toxins produced by *Bacillus thuringiensis* (*Bt*). In the present study aminopeptidase N (APN) was characterized as Cry toxin interacting protein in larval midgut of castor semilooper, *Achaea janata*. Transcriptomic and proteomic analyses revealed the presence of multiple isoforms of APNs (APN1, 2, 4, 6 and 9) which have less than 40% sequence similarity but show the presence of characteristic “GAMENEG” and zinc-binding motifs. Feeding of sublethal dose of Cry toxin caused differential expression of various APN isoform. Further, 6^th^ generation Cry toxin exposed larvae showed reduced expression of APN2. This report suggests that *A. janata* larvae exploit altered expression of APNs to overcome the deleterious effects of Cry toxicity, which might facilitate toxin tolerance in long run.

## Introduction

Proteases are widely analysed proteolytic enzymes found from lower to higher organisms including insects (Reeck et al., 1999). Insects midgut secretary and brush border proteases play a vital role in dietary nutrient metabolism (Terra and Ferreria, 1994; Naseri et al., 2010; Sarate et al., 2012; Jagdale et al., 2017). Among various classes of proteases identified in insects, aminopeptidase N (APNs) are primarily metallo-aminopeptidase with broad specificity. They sequentially and preferentially cleave the neutral amino acids from the N-terminal end of a peptide/protein (Terra and Ferreira, 1994). In addition to the digestive role of APNs, they have been characterized it as Cry toxin recptors/ binding proteins in large number of lepidopteran insects (Jenkins and Dean, 2000) and were shown to mediate *Bacillus thuringiensis* (*Bt)* Cry toxicity (Bravo et al., 2004; Soberan et al., 2009; Adang et al., 2014).

Cry toxins produced by various strains of *Bt* are known to be target specific and have defined spectrum of insecticidal activity against the host insects. This is mediated by the different Cry interacting/binding proteins localized in the apical brush border membrane of the midgut cells, mediated either by pore formation or by signal transduction (Soberon et al., 2009). The pore formation model involves series of events like toxin solubilization, activation of protoxin, its binding to cadherins, removal of one α-helix, oligomerization, interaction with GPI-anchored aminoprptiadse N (APNs)/ Alkaline phosphatises (ALPs)/ ATP-binding cassette transporters (ABBC2), membrane insertion, pore formation leading to cell damage (Bravo et al., 2004, Crickmore, 2005; Guo et al., 2015; Soberón et al., 2018).While in signal transduction model, *in vitro* studies demonstrated the involvement of cAMP dependent prtoein kinase A (PKA) pathway (Zhang et al., 2006). However, in CF1 cells the Cry toxicity was not associated with PKA pathway (Portugal et al., 2017). Recent studies revealed that Cry toxin induces necrotic cell death in *B.mori* ABCC2 transfected Sf9 cells (Endo et al., 2017) as well as in larval midgut of *A. janta* (Chauhan et al., 2017).

Aminopeptidase N (EC.3.4.11.2) has been identified as Cry toxin receptor in various Lepidopteran insects like *M. sexta* (Knight et al., 1995), *B. mori* (Yaoi et al., 1998), *Lymantria dispar* (Garner et al.,1999), *Plutella xylostella* (Nakanishi, et al., 2002) *Spodoptera litura* (Agrawal, et al., 2002) *H. virescens* (Banks, et al., 2003) and *H. armigera* (Rajagopal et al., 2003). These are a class of peptides belonging to Zn^++^ dependent gluzincins family of M1 type metalloproteases (Hooper, 1994; Albiston et al., 2004). APNs show special features like aminopeptidase motif “GAMENWG”, Zn^++^ binding motif “HEXXHX_18_E”, N-terminal with the signal peptide and C-terminal with GPI-anchor peptide, which facilitates their attachment with brush border. *In silico* analysis of sequences also revealed the presence of many O and N-glycosylation sites which were predicted to facilitate Cry toxin binding (Pigott and Ellar, 2007). In lepidopteran insects, APNs are classified into eight clusters (APN1-8) based on their amino acid sequence identity/similarity (Crava et al., 2010; Hughes, 2014; Lin et al., 2014). Multiple isoforms of APNs to be expressed in the midgut of *H. armigera* (Angelucci et al., 2008), *Ostrinia nubilalis* (Crava et al., 2010; 2013; Khajuria et al., 2011; *Epiphyas postvittan* (Simpson, et al., 2008) *B. mori* (Nakanishi, et al., 2002; Crava et al., 2010) and *Trichoplusia ni* (Wang et al., 2005; Tiewsiri and Wang, 2011). Further in lepidopteran genome these isoforms were organized in a single cluster. (Crava, et al., 2010; d’Alencon et al., 2010; Baxter et al., 2011; Gao et al., 2019). However, uncertainty about the exact isoform which act as functional receptor in a given species (Bretschneider et al., 2016; Zhang et al., 2019).

Bt-based biopesticides are currently being used as an alternative to chemical insecticides for the effective management of economically important Lepidopteran pests either Bt based formulations or Bt transgenic plants. However, development of resistance against Cry toxin based control strategies in insect populations is a great concern and needs a careful evaluation (Tabashniket al., 2013). Reports related to the development of resistance in laboratory based as well as field collected populations shown to mediated by multiple mechanisms (Pardo-Lopez et al., 2013; de Bortoli and Jurat-Fuentes, 2019). In certain species, it was due to the alteration in Cry toxin activation by gut proteases (Keller et al., 1996; Oppert et al., 1997; Li et al., 2004), sequestration of Cry toxins by glycolipid moieties (Ma et al., 2005) and an elevated immune response (Hernández-Martínez et al., 2010) were additional mechanisms which promoted the development of Cry resistance. Mutation in APN receptor proteins was shown to be associated with reduced interaction of Cry toxin with larval midgut (Zhang et al., 2009). Tiewsiri and Wang (2011) correlated toxin resistance with a reduced and differential expression of various APN isoforms in *T. ni*. Recently Sun et al., (2019) demonstrated that knockdown of APN isoforms 6 and 8 decreases the susceptibility of *Chilo suppressalis* larvae towards Cry toxins.

In the present study, an attempt was made to characterize the aminopeptidase N as Cry toxin receptor/binding protein in the larval midgut of *Achaea janata*, commonly known as castor semilooper a serious lepidopteran pest of castor plantation in India. Further, various isoform of APNs were identified and their expression profile during continuous sublethal toxin exposure as during generation wise feeding of sublethal toxin in each successive generation for six generations was analyzed. The results obtained clearly revealed altered expression of various APN isoforms during long term generation wise sublethal Cry toxin exposure. The finding of the present study suggests that this altered expression of APN isoforms might be responsible to overcome the toxicity towards Bt formulations (DOR Bt-1) presently using for the management of castor semilooper in local castor fields.

## Material and Methods

### Insect culture maintenance

Egg masses were collected from the agricultural fields of Telangana region; India where the crops were never sprayed with *Bt*-based biopesticides. In the laboratory immediately after hatching, the neonates were transferred onto fresh sterile castor leaves. The larvae were maintained in insect culture facility under of 14:10 h (light: dark period), 70±5% RH and 27±2°C temperature until pupation. The glass troughs containing sterile moist sand were used for maintaining the pupae which were allowed to emerge into adults. The adult moths (male and female) were collected and transferred to breeding cages, where they were fed with 10% honey (Pavani et al., 2015). Adult female moths laid eggs on fresh castor leaves surface, which hatched in 4-5 days. In the laboratory, three continuous generations were maintained to get homogenous population. Later based on the experimental requirement the larvae were exposed to sub-lethal dosage of Cry toxin treatments and the control cultures were simultaneously maintained throughout the study.

### Tissue isolation

The 3^rd^ instar larvae were narcotized by placing them on ice for 15-20 min. The midgut tissue was isolated according to the procedure described in Chauhan et al, (2017). The isolated midguts were used for the isolation of total RNA and brush border membrane vesicles (BBMVs) preparation.

### Preparation of midgut brush border membrane vesicle (BBMV) fractions

The midgut membrane fraction was prepared using the protocol of Wolfersberger et al, (1987) with slight modifications. Dissected midgut tissue was transferred to centrifuge tube containing ice-cold MET buffer (300 mM mannitol, 17 mM Tris-HCl pH 7.5, 5 mM EGTA and 1 mM PMSF), vigorously vortexed and briefly centrifuged for 5 min at 1000 X g to obtain plant debris free clean tissue. Homogenate was prepared using ice-cold MET buffer (10% w/v) with glass homogenizer. The resultant homogenate was mixed with equal volume of MgCl_2_ (24 mM), by vigorous shaking and place on ice for 15 min. It was centrifuged at 4°C (2500 X g) for 15 min. The supernatant was further centrifuged at 30,000 X g for 1 h at 4°C. The resulting pellet which contained BBMV’s fraction was re-suspended in HEPES buffered saline (10 mM HEPES, pH 7.4, and 150 mM NaCl). The purity of the obtained BBMV’s was checked by alkaline phosphatase activity assay (Wolfersberger et al., 1987), and the protein concentration was estimated using method of Brodford, (1976). Pure BBMVs fraction was flash-frozen in liquid nitrogen and stored at −80° C until use.

### Ligand blot analysis with midgut BBMV proteins

The Cry crystals obtained from DOR *Bt*-1 and Cry1Ac toxin (isolated from recombinant clone obtained from Bacillus Genetic Stock Centre, Ohio, USA) was solubilised in alkaline pH (Na_2_ CO_3_. NaH CO_3_ buffer pH 9.5), and the protoxin was activated by trypsinization (toxin: bovine trypsin; 50:1 (w/w)) at 37^0^C for 2h for getting activated Cry toxin (Osir et al., 1999). The midgut BBMV proteins (10 μg) were separated by 10% SDS-PAGE and electrotransferred to a nitrocellulose membrane. The membrane was blocked using blocking buffer [5% (w/v) skimmed milk powder in 0.01 M Tris buffer saline (TBS, pH 7.4)] for 1 h, followed by incubation in 5 ml of blocking buffer containing of activated DOR *Bt*-1 or Cry1Ac toxin (1μg/ml) for 1 h at room temperature. Following this blot was individually incubated with polyclonal primary antibody generated against DOR *Bt*-1 toxins or anti-Cry 1Ac toxin antibody (1:500 dilution; both raised in mice) for overnight. It was then incubated with anti-mouse HRP conjugated secondary antibody (Thermo Fisher Scientific, USA). Finally, blot was developed using chemiluminescence kit (Takara Bio Inc, Japan).

### APN and Cry toxin(s) interaction analysis

For this midgut was fixed in Bouin’s fixative (saturated picric acid: formaldehyde: glacial acetic acid; 15:5:1), processed and embedded in paraplast (Sigma-Aldrich, USA). The transverse sections (~5 μm) were prepared using a rotary microtome (Leica Microsystems, Germany). The slides were deparaffinised using xylene and hydrated through decreasing ethanol series. For immunostaining, sections were washed with PBS followed by PBST, then blocked using 10% goat serum (Thermo Fisher Scientific, USA). 1 μg/ml of Cry toxin (Cry1Ac or DOR *Bt*-1) was added on to the slides and incubated for 2 h at room temperature. Primary polyclonal antibody used was generated against conserved aminopeptidase N sequence (1:500 dilution; raised in rabbit) and anti-Cry toxin(s) antibody (1:500 dilution; generated in mice) were added and the slides were incubated overnight at 4°C. The slides were further washed with PBS solution to remove unbound primary antibodies then respective secondary antibody (anti-rabbit goat IgG-FITC conjugated (green) and anti-mice goat IgG-TRITC (red)) were added to the sections and incubated at RT for 2 h. The excess of antibody was washed with PBST and the slides were mounted using gold antifade mountant (Thermo Fisher Scientific, USA) for prolonged storage. The sections were visualized under fluorescence microscope. (Leica Microsystems, Germany).

### Generation of toxin exposed insect population

DOR *Bt*-1 formulations (based on a local strain of *B. thuringiensis*) obtained from Indian Institute of Oilseeds Research, Hyderabad which has potential insecticidal toxicity towards larval forms of *A. janata*. The advantage of this stain is it has both *cry1* (*cry 1Aa, cry 1Ab* and *cry 1Ac*) and *cry2* (*cry 2a*, and *cry 2b*) genes (Reddy et al., 2012). The LC50 reported for this formulation for 3^rd^ instar larvae is 247.52 μg/ml (Vimala devi et al., 2006). For the present study sub-lethal dosage of formulations used was 0.170 μg/cm^2^ (Chauhan et al., 2017). Fresh castor leaves were coated with it and 3^rd^ instar larvae were released. The insect midgut samples from susceptible larvae were collected at every 12 h time interval of continuous toxin exposure. For a generation of toxin exposed insect population, the 3^rd^ instar larvae were exposed for 3 days on sub-lethal toxin coated leaves, followed by their transfer on normal leaves for the recovery (3 days). Once again the larvae were exposed to toxin coated leaves for 3 days. Finally, they were maintained on normal leaves to complete their life cycle. The surviving insects were maintained for the next generation. At each generation, the required tissue samples were collected from 3^rd^ instar larvae (prior to toxin exposure) and experiments were carried out. Using the above protocol toxin exposed population was generated and present analysis was carried out up to the sixth generation.

### RNA isolation and cDNA preparation

The midgut tissue dissected out from continuous toxin-exposed susceptible and toxin exposed insects for 6 generations (n=5) were processed for RNA isolation using TRI reagent TM (Sigma Aldrich, USA) following the manufacturer’s instructions. Quantity and quality of isolated RNA were checked using Nanodrop 1000 (Thermo Scientific, USA) and integrity of RNA was assessed by performing denaturing electrophoresis on a 1% formaldehyde agarose gel (Green and Sambrook, 2012). First strand cDNA synthesis was carried out according to manufacturer’s protocol (Invitrogen, Life Technologies, USA) by taking the 1μg of total RNA.

### Cloning, characterization and expression analysis of APN isoforms

As the genomic sequence of *A. janata* is not available in database hence an attempt was made to carry out next generation RNA sequencing (*denovo* transcriptome analysis) of larval midgut (Sequence Read Archive (SRA) No. SRR7212215)). Based on the homology search in NCBI database, five different isoforms of APNs were identified. Using standard protocol, full length sequences of the various isoforms were cloned using ORF-specific primers (Table 1). Detailed *in silico* analysis of these putative APNs was carried using bioinformatics tools (multi alignment, homology and conserved domain analyses). Further the mRNA expression (transcript) was analyzed upon either continuous sub-lethal toxin exposure or in toxin tolerant insect population using qRT-PCR with gene primers (Table 1). For quantitative PCR analysis, a 20 μl reaction was carried out using a SYBR Green Real-Time PCR Master Mix (Thermo fisher Scientific, USA). Insect rS7 (KF984201) was used as an internal reference to normalize the transcript expression levels (Chauhan et al., 2017).The real-time expression analysis was performed in triplicates. Relative expression of various genes was employing method of Livak and Schmittgen (2001).

**Table 1:**
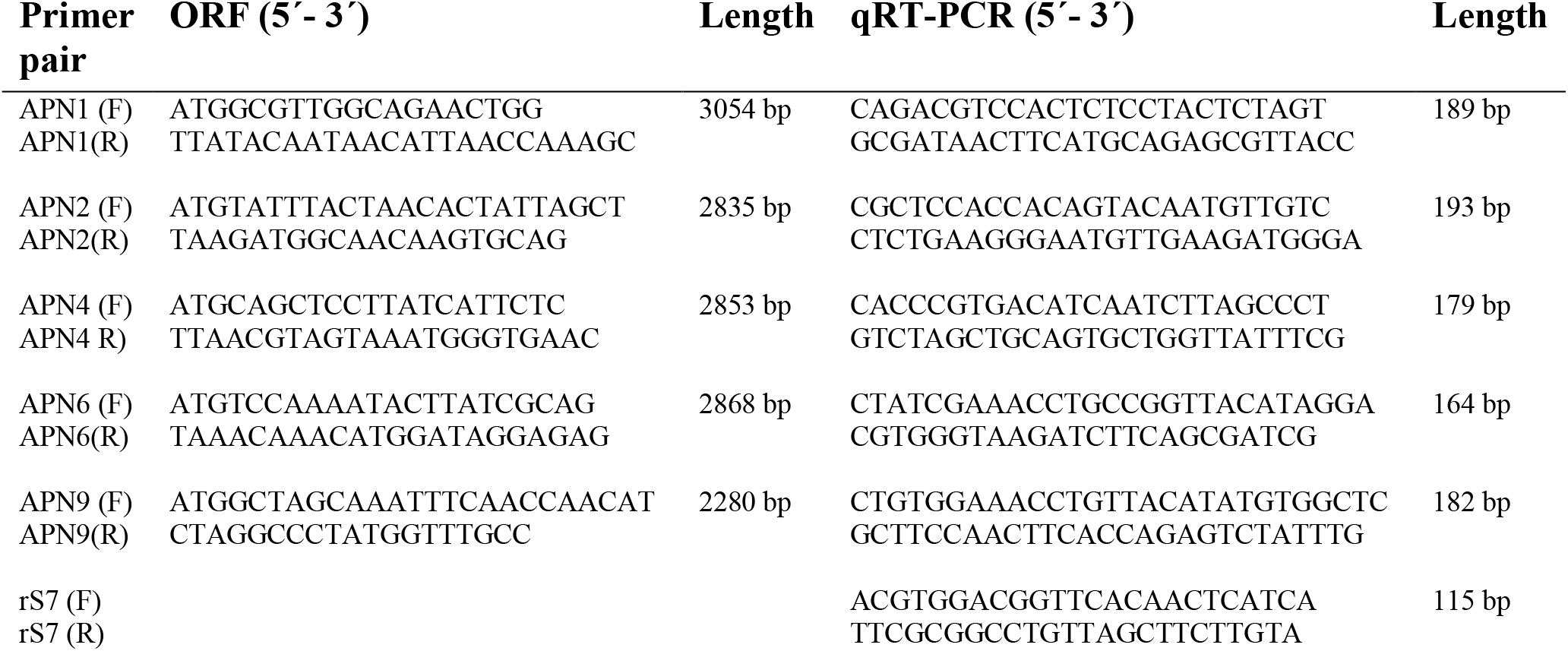
List of primers used for cloning of full length (ORF) of APN isoforms and Real time primers used for the expression analysis in toxin exposed and toxin tolerant insect population.

### Two-dimensional electrophoresis and MALDI-MS-MS analysis

The midgut BBMV proteins were subjected to clean-up using the 2-D Clean-up Kit (GE Healthcare Life Sciences, USA) as per the manufacturer’s instructions. Immobiline Dry strips (4-7 pH 7 cm) were rehydrated overnight in 125 μl of rehydration buffer (7.0 M urea, 2.0 M thiourea, 4% CHAPS, 50 mM DTT and 1% IPG buffer) containing 100 μg of protein. The rehydrated IPG strips containing 100 μg of proteins were then subjected to isoelectric focusing as per the manufacturer’s standard protocol (GE Healthcare, USA). After IEF, equilibrated for 20 min in solution (6 M urea, 30% glycerol and 2% SDS in 0.1 M Tris-HCl, pH 8.8) containing 50 mM DTT followed by 100 mM IDA and second dimensional SDS-PAGE was carried out in duplicates. After electrophoresis one gel was used for immunoblotting with APN antibody and a second one was stained using Coomassie brilliant blue R-250. For MALDI-TOF-TOF analysis of excised protein spot from 2-D gel was subjected DTT reduction followed by alkylation before trypsin digestion. The peptide mass fingerprint (PMF) spectrum of peptides mixture was generated (MALDI-MS) and high intensity PMF peaks were further ionized (MALDI-MS-MS) for *de novo* sequencing. Then, PMF spectrum of 2-D protein spot was matched with theoretically digested APN peptide mass fingerprint due to non-availability of available reports on *A. janata* protein data report.

### Differential two-dimensional gel electrophoresis (2D-DIGE)

The midgut BBMV samples were prepared from GC6 (generation 6 control) and GT6 (sixth generation larvae obtained after a sub-lethal dosage of Cry toxin exposure during every generation). Samples were cleaned up using 2-D Clean-up kit (GE Healthcare Life Sciences, USA), the obtained pellet was dissolved in 2-D lysis buffer containing 7 M urea, 2 M thiourea and 4% CHAPS without dithiothreitol (DTT). The protein quantification was done using Ettan 2-D Quantification kit (Amersham Biosciences, USA). Supplied CyDyes (Cy3/Cy5/Cy2 (5 nmol each)) (GE Healthcare, UK) was reconstituted using freshly opened 99.8% anhydrous dimethylformamide (DMF) (Sigma Aldrich, USA) to a working solution (400 pmol) for protein labelling. The pH of the protein solution was adjusted between 8.0-8.5 with 0.5 N NaOH before protein labelling with dyes. Labelling of BBMV protein samples were prepared from control (G0) as well as Cry toxin-exposed larvae (G6) was done with 400 pmol of Cy3 and Cy5 dyes respectively as per the supplier’s instructions. An equal amount of protein from all the twelve samples [G0 (6) + G6 (6)] were mixed for generating internal standard, which was labelled with Cy2 and equally distributed to all. This facilitated in generating accurate spot statistics between gels and increased the level of confidence. After incubation in the dark for 30 min on ice, the dye quenching was achieved by addition of 10 mM L-lysine for 10-20 min in dark. Six biological replicates were used for this experiment; dye swapping was done to minimize the artefacts generated by the dye components.

Each gel consisted of G0 and G6, samples (50 μg, each) labelled with Cy3 or Cy5 and internal standard labelled with Cy2. Finally, the required volume (340 μl) was made with 2-D rehydration buffer containing 40 mM DTT and 0.5% IPG buffer pH 4-7. After rehydration for 12-16 h, IPG strips (pH 4-7, 18 cm) were subjected to isoelectric focusing (IEF) in EttanIPGPhor 3 system as per standard protocol. Equilibration of IPG strips was done with buffer containing 6 M urea, 30% glycerol, 2% SDS and 50 mM DTT for 20 min followed by alkylation with 100 mM iodoacetate (IDA) (Sigma Aldrich, USA) for 20 min. Finally, IPG strips were equilibrated in SDS-PAGE cathode buffer, before performing the second dimension analysis. After electrophoresis, gels were scanned at different excitation and emission wavelength for Cy3, Cy5 and Cy2 labelled proteins using Typhoon variable mode scanner (Amersham Biosciences, USA). All the gels were scanned with PMT of 600 and 200 μm/pixel resolutions. Images were processed using DeCyder 2-D version 7.0 software through Differential In-gel Analysis (DIA) and Biological Variation Analysis (BVA). Finally, differentially regulated protein spots were detected with a significant p-value of <0.05 generated by the student t-test and one-way ANOVA.

### *In situ* localization of APN2

*A. janata* APN2 cRNA probes were prepared using T7 promoter sequence (5’-<u>TAATACGACTCACTATAGGG</u>CGCTCCACCACAGTACAATGTTGTC-3’) based primer was used for synthesis of the antisense probe (positive), while Sp6 based promoter sequence(5’-<u>ATTTAGGTGACACTATAG</u>CTCTGAAGGGAATGTTGAAGATGGA-3’) was used for sense probe (negative). *In vitro* transcription of DNA template was carried out using thermocycler followed by RNA probe synthesis using DIG RNA labelling kit using manufactures protocol (Roche Diagnostics, Germany). DIG labelled mRNA was synthesized and analysed using agarose gel electrophoresis and sequencing.

Cryotome cut midgut transverse section was fixed on the clean glass slide and washed with PBST buffer. Then it was permeabilized using proteinase K (1 μg/ml) and fixed again with 4% PFA. After fixation the slides were incubated with 200 μl hybridization buffer (50% formamide, 5X SSC, 1% SDS, 5 mM EDTA, 0.1% CHAPS, and 50 μg/mlheparin) for 1 h at 50°C. Laboratory prepared DIG labelled cRNA (added with 200 μl of hybridization buffer) probe was used and heat denatured by incubating it for 5 min at 80°C.While incubating the slides were covered with parafilm and kept in sterile incubator. After cRNA hybridisation, the slides were washed using wash buffer (SSC, 50% formamide, 0.1% Tween 20), five times for 5 min at 50°C. Each time SSC concentration was gradually decreased followed by incubation of slides with solution A (10 Mm Tris-HCl pH 8.0, 0.5 M NaCl, 5 mM EDTA, containing 0.1% Tween 20) at RT for 5 min. To avoid any irrelevant contamination reaction was supplemented with RNase A (20 μg/ml) and incubated for 20 min at RT. After RNase treatment the probed sections slides were washed twice for 5 min with solution A. Followed by washing with maleic acid buffer (150 mMNaCl, 100 mM maleic acid, 0.1% Tween 20, pH 7.5) for enhanced antibody reaction at RT for 5 min, incubated with 5% BSA blocking solution at RT for 1.5 h followed by incubation with anti-Digoxigenin-FITC antibody [21H8] (FITC) (ab119349) (1: 500) at 4°C for overnight. These slides were washed four times for 15 min with DIG wash buffer (Roche Diagnostics, Germany). Visualization of fluorescent labelled section was carried out using fluorescent microscope at 480 nm wavelength (Leica Microsystems, Germany).

### Statistical analysis

The validation of the data was carried out using SEM (standard error of mean) using Three biological replicates (n=3). It was also proceeds for homogeneity and normality tests. Sigma Plot v12.3 software were used for the verification of Statistical significance between the compared values. One-Way ANOVA was used for calculating the significance between the tested groups. This was followed by Student-Newman-Keuls (SNK) test for pair wise multiple analysis. For each set up experiment p-value was calculated.

## Results

### Characterization of aminopeptidase N as midgut Cry toxin receptor

SDS-PAGE separated midgut BBMV proteins were electrotransferred on nitrocellulose membrane. Ligand binding analysis (toxin overlay assay) was carried out using both Cry1Ac and DOR *Bt*-1toxin(s). The results presented in figure 1 clearly show that Cry toxin(s) interacted with more than 5 gut membrane proteins. Among them the prominent interaction was seen with ~110 kDa protein, with Cry1Ac (Fig 1, L3) as well as DOR *Bt*-1 toxins (Fig 1, L4). Immunoblotting studies using APN antibody, revealed strong immuno cross reactivity with ~110 kDa protein, suggesting that it was an APN (Fig 1, L6). Further the peptide sequence “AQIVNDVFQFARS” obtained through MALDI-TOF/TOF analyses of this protein band matched with the alanyl aminopeptidase of *H. virescens* with molecular weight ~113 kDa (Gill et al., 1995).

**Figure 1.**
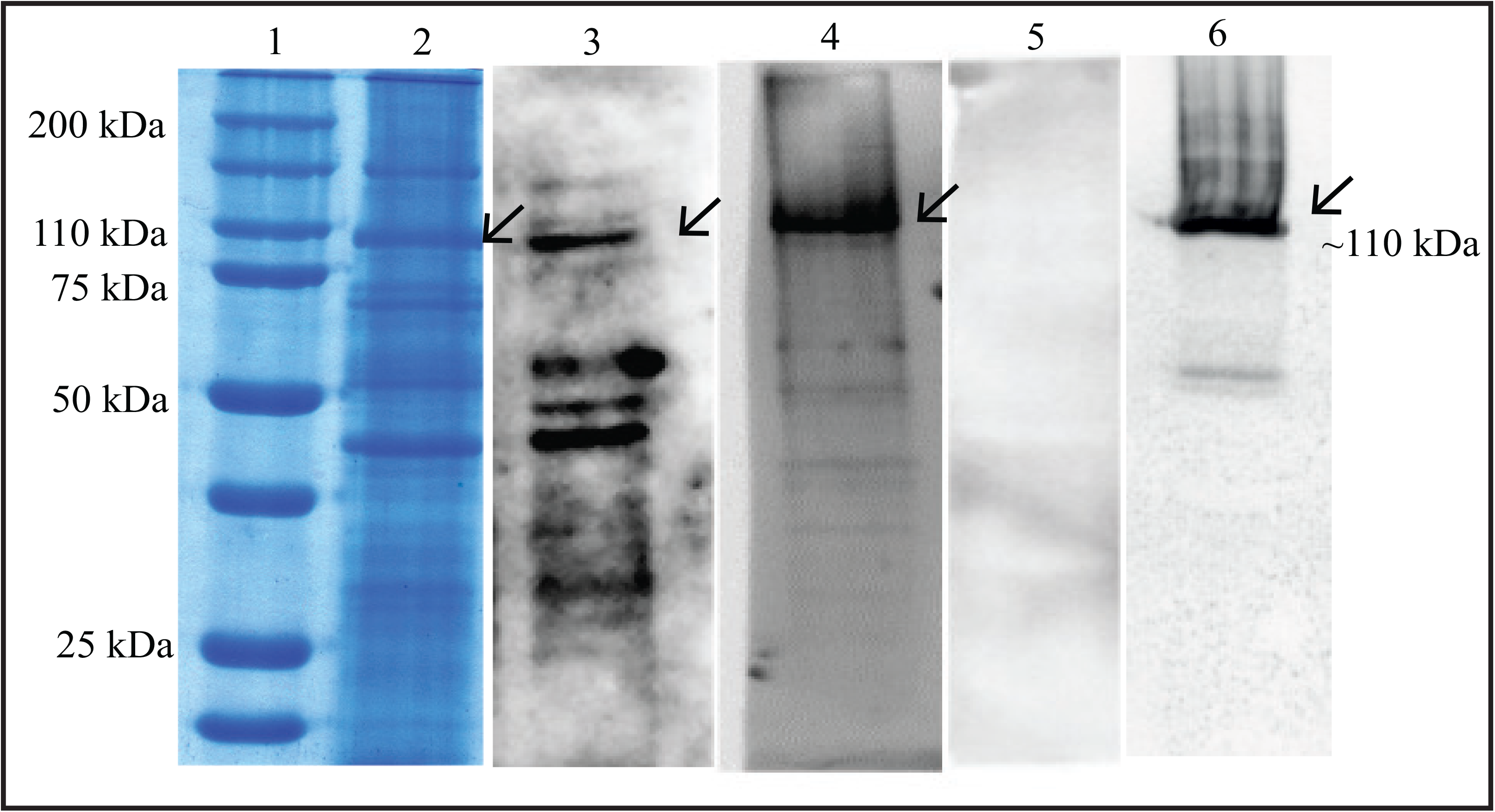
Mid-gut membrane protein and Cry toxin interaction analysis. SDS-PAGE separation of BBMV proteins isolated from *A. janata* larval mid-gut (Lane 2). Lignad blot analysis (toxin overlay assay) with Cry1Ac (Lane 3) and DOR *Bt*-1 (Lane 4) toxins show multiple toxin interacting membrane proteins and the prominent interaction was seen with ~110 kDa protein (←). Lane 6 shows immunolotting with APN antibody (←). Lane 1 is protein molecular weight marker. Lane 5 shows ligand blot with BSA (negative control) (Complete western blot image is provided in supplementary Fig. 3).

The presence of APN in inner apical brush border membrane of midgut epithelium was visualized through immunofluorescence using APN antibody (Fig 2A, b). Colocalization studies showed presence of APN as well as Cry toxin(s) signal in inner brush border membrane of midgut epithelial cells in close proximity (Fig 2B, Panel C). The signal was visible in the Cry1Ac (Fig 2B, c1) as well as DOR *Bt*-1 (Fig 2B, c2) toxin incubated sections but not with BSA (Fig 2B, c3). These findings clearly suggested that the interactions were specific and Cry toxins selectively bound to the brush border membrane proteins, where APNs are known to be localized.

**Figure 2.**
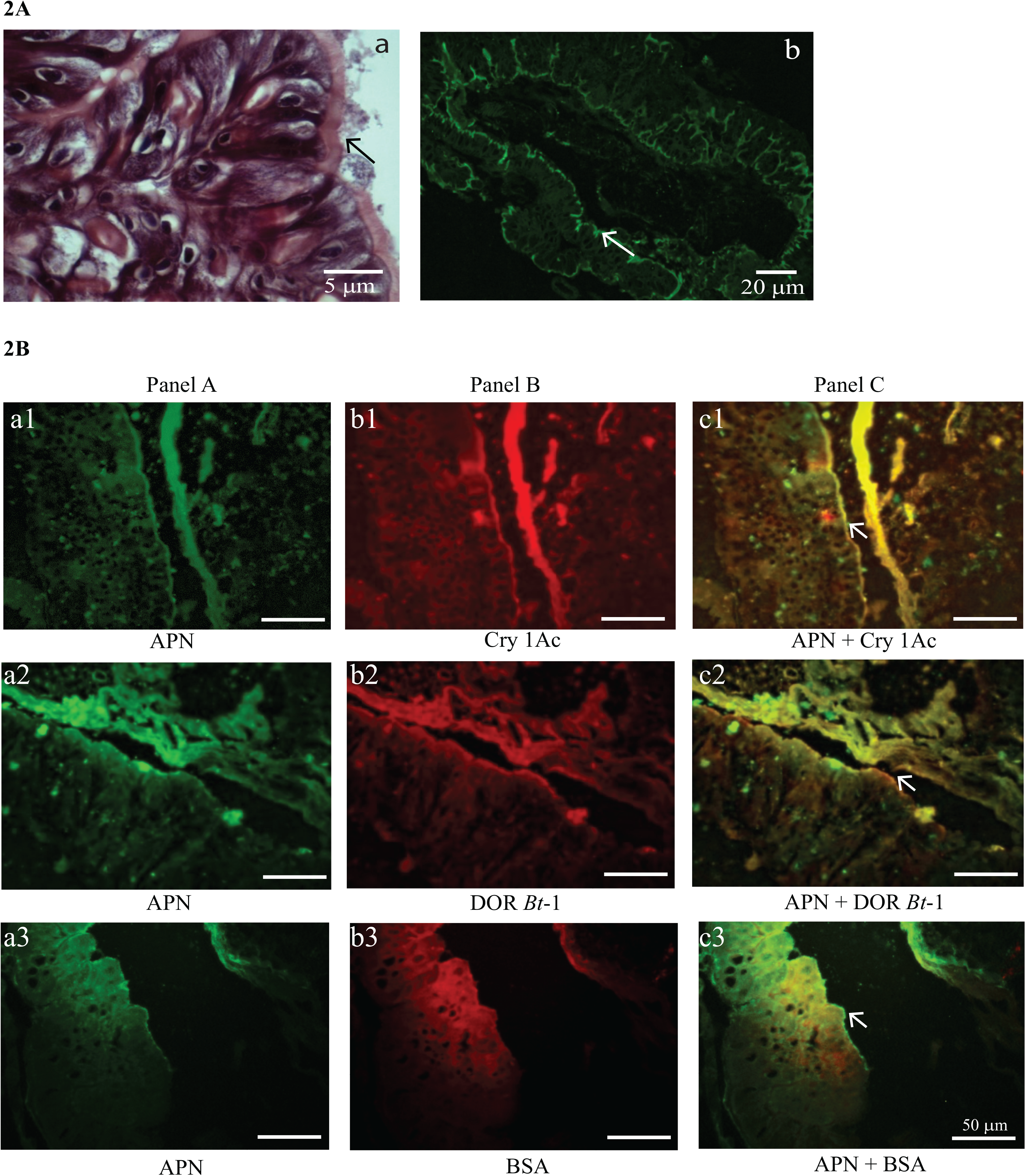
Immunofluorescence based detection of Aminopeptidase N and Cry toxin in *A. janata* mid-gut cross-section. (2A) Micrograph showing cross section of larval mid-gut. (a) Stained with haematoxylin/eosin, note the presence of apical brush border of mid-gut epithelium (←). (b) Localization of APNs in epithelial brush border (dark green) using immunofluorescence with APN antibody. (2B) Aminopeptidase N and Cry toxin interaction analysis using immunofluorescence. Panel A (a1, a2 and a3) shows the detection of APN on apical brush border of mid-gut epithelium using APN antibody. Panel B shows the interaction of Cry toxin (b1: Cry1Ac; b2 DOR *Bt*-1) with the apical brush border. Panel C shows the colocalization of both APN and Cry binding regions. Please note that merged image shows the presence of both the molecules APN and Cry toxin in the same region of the apical membrane of mid-gut (c1 and c2; ←). BSA was used as negative control where no signal was detected with Cry antibodies at the brush border (b3 and c3), which is clearly seen in c1 and c2

### APN isoform analysis

Full length of cDNA for all the five isoforms of APN detected in *A. janata* midgut larval transcriptome was cloned into pTZ57R/T vector (Thermo Scientific, USA) and Sanger sequenced. The sequences were confirmed and submitted to Gene Bank (https://www.ncbi.nlm.nih.gov/BankIt/) with accession numbers - APN1 (GenBank accession no. **<u>MF425653</u>**), APN2 (GenBank accession no. **<u>MF425654</u>**), APN4 (GenBank accession no. **<u>DQ872666</u>**), APN6 (GenBank accession no. **<u>MF425655</u>**) and APN9 (GenBank accession no. **<u>MF425656</u>**). Deduced translated amino acid sequences of cloned isoforms of *A. janata* were aligned through Multiple Sequence Alignment (MSA) (http://www.ebi.ac.uk/Tools/msa/) and the search for the presence of characteristic aminopeptidase activity motif “GAMENEG” and zinc-binding motif “HEXXHX_18_H” showed the presence of both of them (S1a Fig). Similarity between the isoforms was checked using Percent Identity Matrix (Clustal 2.1) and the result revealed that the cloned isoforms had less than 37% sequence identity (S1b Fig). Furthermore, conserved domain analysis of these sequences revealed that they belong to “Peptidase M1 aminopeptidase N” family with ERAP1 with M1_APN_2 and ERAP1_C domains. Transmembrane peptide was found to be present at C-terminal end of APN1, APN2 and APN4 while it was absent in APN6 and APN9. Further APN1, APN4 and APN6 possessed signal peptide at N-terminal, which was absent in APN2 and APN9. It is interesting to note that APN9, lacked both N and C-terminal signal sequences (S1c Fig).

### Differential expression of APN isoforms in toxin exposed larval populations

In susceptible insect larvae, the qRT-PCR analyses revealed the down-regulation of all the midgut APN isoforms in time-dependent manner from 12 to 72 h upon continuous exposure to sub-lethal dosages of Cry toxin(s) (Fig 3). Further, the isoforms were differentially regulated and the expression of each isoform differed significantly from one another during the various time points of Cry toxin exposure (Fig 3).

**Figure 3.**
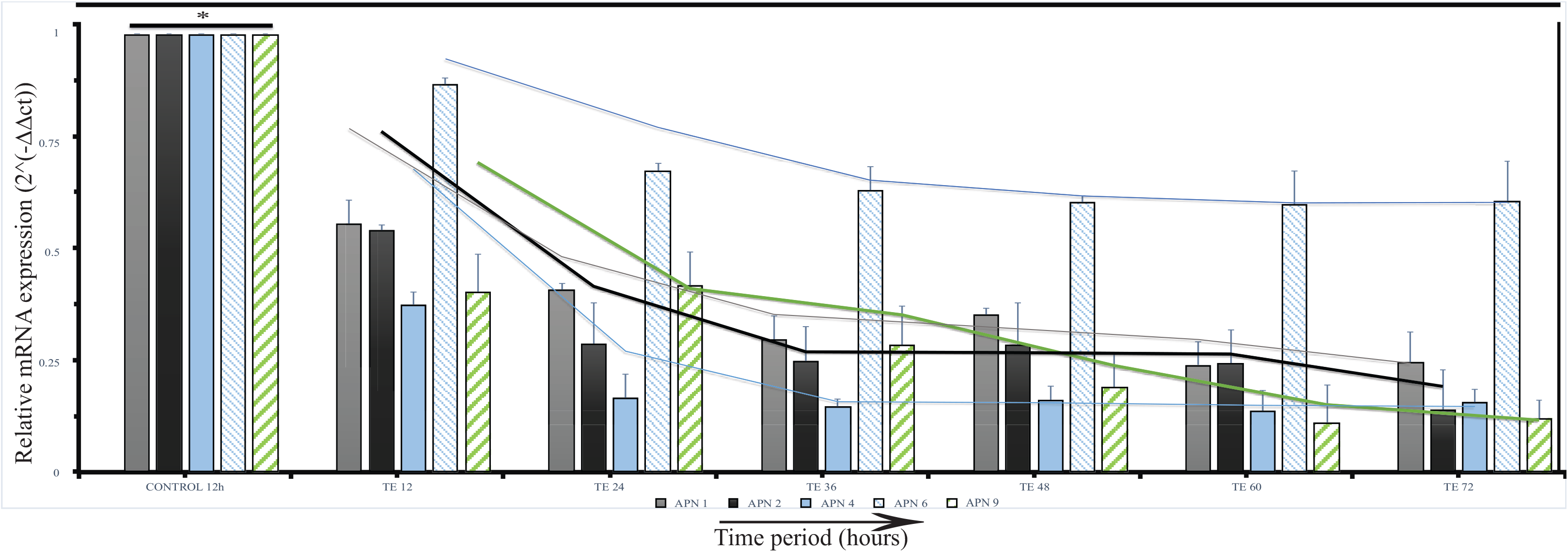
Expression analysis of mid-gut aminopeptidase N isoforms using qRT-PCR during sub-lethal toxin exposure. A gradual decline in expression of various isoforms from 12 to 72 h of Cry intoxication. Please note the differential down regulation of each isoform in various time points. *p < 0.005 significance between experimental groups (n=3).

The generation wise expression analyses, form 6 generation toxin-exposed insects revealed that the expression of isoforms APN1, APN2, APN4 and APN 6 was up-regulated in G1, as compared to control (G0) (Fig 4). On the other hand expression of APN9 was low in G1 and G2 generations and increased gradually from G3 to G6 and reached more or less at level of G0. It is interesting to note that expression of APN 2 declined gradually from G2 and G3 and remained at low level till G6. APN 6 also showed lower expression similar to APN2 up to G3 then it increased marginally from G4 to G6. However, high APN4 expression seen in G1 declined gradually up to G3 but increased significantly from G4 to G6 generation, where it showed higher expression as compared to G0. The results presented above clearly showed that in generation 6 (G6) there was a reduced expression of APN2, while expression of APN1 and APN6 was slightly higher. It is interesting to note that APN4 expression was significantly high in G6, when compared with control generation G0 (Fig 4). A comparison in the expression profile showed down-regulation of APN2 and up-regulation of APN4 in generation wise analyses (Fig 4f).

**Figure 4.**
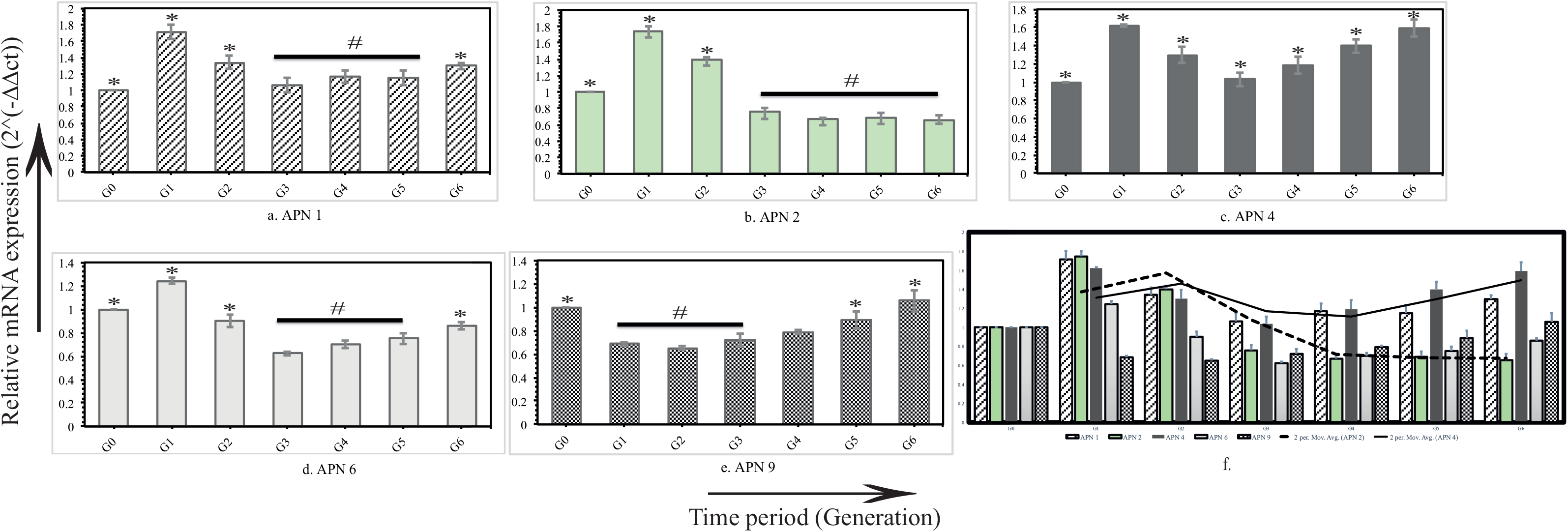
Profiling of APN isoforms in toxin exposed insect population. All the APN isoforms (APN1, APN2, APN4, APN6 and APN9) showed significant up regulation in G1 generation when compared with G0 (a, b, c and d) except APN9 (e). Please note the expression of various APNs declined gradually in G2 and G3 (a, b, c, d and e) except APN2 (b) once again up regulated from G4 to G6 (a, c, d and e). However it is interesting to note that the expression of APN2 not only gradually declined but also remained fairly low from G3 to G6 generations (b and f), while expression of APN4 was significantly increased from G3 to G6 generation (c and f) (*p < 0.005 significance between experimental groups (n=3) and # represent non-significant to each other).

### Proteomic analysis of APN isoforms

Immunoblot analysis using anti-APN antibody showed its interaction with multiple protein spots (~9spots) in two-dimensional gel electrophoretic gel (S2 Fig). This clearly indicated the presence of multiple isoforms of aminopeptidase in the larval midgut of *A. janata*. Using DIGE protocol midgut proteins from G0 (control larval populations) and G6 (toxin exposed larval population) were analysed. The results showed a total of 1552 spots in which 27 protein spots were found to be differentially expressed. For statistical analysis, a cut off of ± 2.0 (average ratio) was used. Then student t-test (>0.05) and one way ANOVA (>0.05) was performed. For the present study, we focused on the spots which interacted with APN antibody (S2 Fig). The differential expression analysis (Fig 5) showed up-regulation (non-significant) of most of the protein spots, except spot number 5 which was moderately down-regulated (Table 2). The spot number 5 was further analysed through MALDI-TOF/TOF and identified as aminopeptidase 2 (APN2) by comparing the theoretical values generated through the same parameters used for protein identification through MS/MS analysis using through Mascot MS/MS ion search online tool (www.matrixscience.com) (Fig 6).

**Figure 5.**
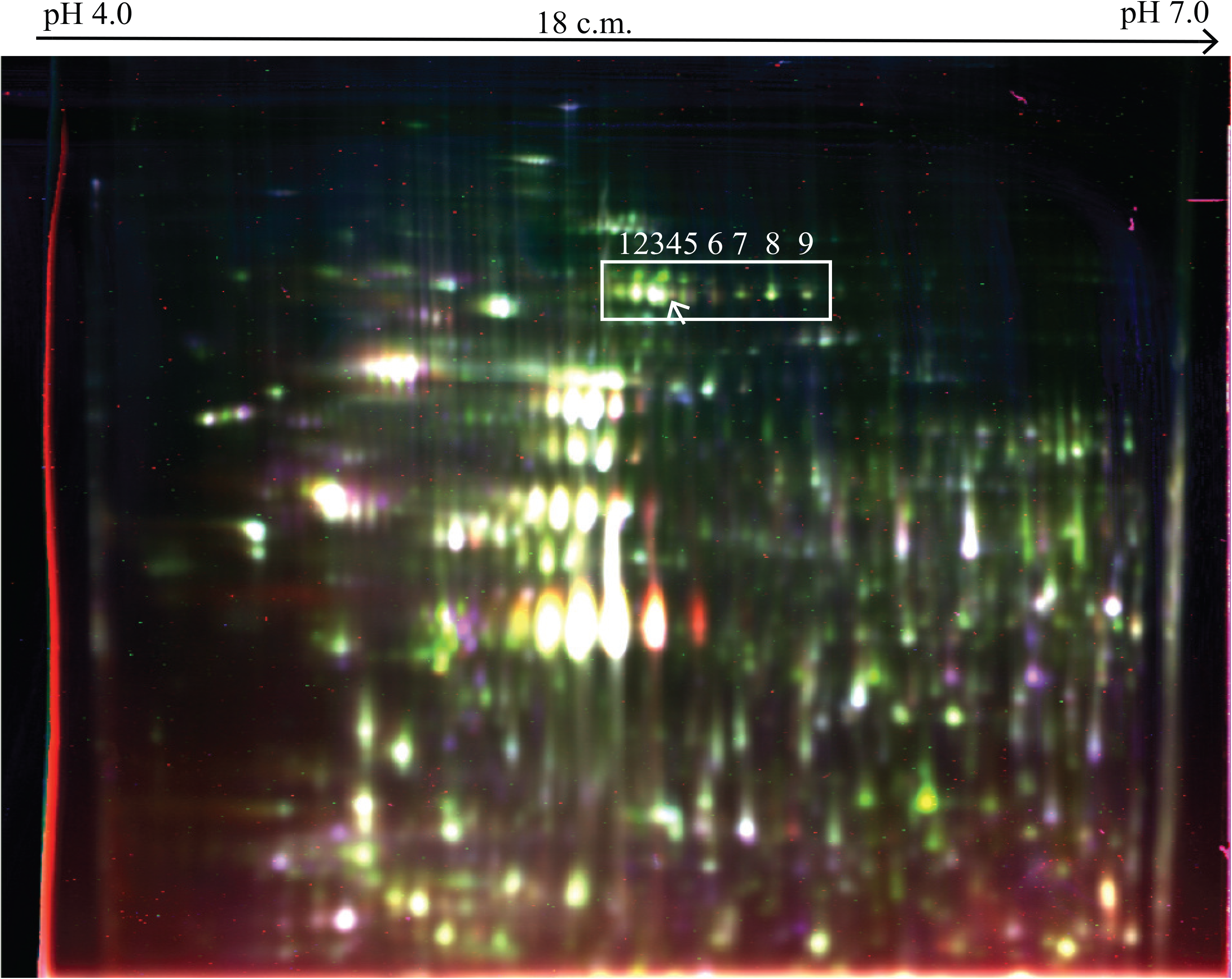
2D-DIGE analysis of *A. janata* mid-gut membrane proteome of G0 and G6 samples. A representative image of membrane protein profile generated by DeCyder software. The result was obtained from experiments conducted with three biological and technical replicates including DIGE swapping to minimise the errors. The series of protein spots indicated in box were identified as aminopeptidase isoforms, using APN antibody (S2 Fig).

**Figure 6.**
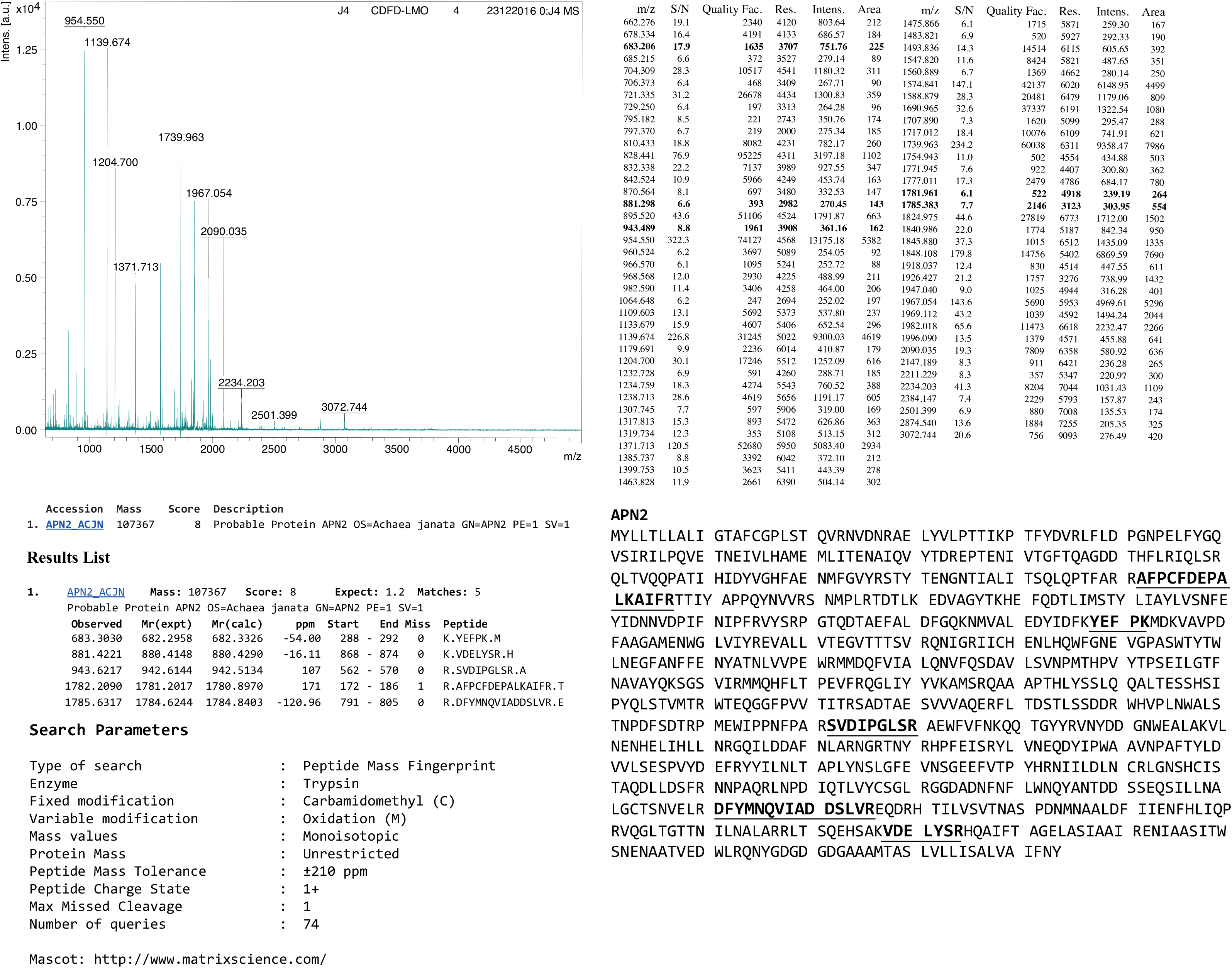
MS/MS analysis of down regulated protein spot using Mascot software. The M/Z value (bold) matched with theoretical peptides generated with APN2 sequence.

**Table 2.** DIGE expression analysis of APN isoforms in G0 and G6 generation which were detected using APN antibody. 2D-DIGE result was analysed using DeCyder 7.0 software provided by GE Healthcare. *Significant (p<0.05), #Not significant.

| Spot<br>No | Master<br>ID | Appearance | T-test | AV. Ratio | One-way<br>ANOVA | Expression<br>Profile |
| --- | --- | --- | --- | --- | --- | --- |
| 1 | 405 | 15/18 | 0.061 | 1.66 | 0.61 | UP * |
| 2 | 416 | 15/18 | 0.33 | 1.41 | 0.33 | UP # |
| 3 | 418 | 18/18 | 0.41 | 1.26 | 0.41 | UP # |
| 4 | 428 | 18/18 | 0.58 | 1.17 | 0.58 | UP # |
| 5 | 425 | 18/18 | 0.56 | -1.07 | 0.56 | DOWN # |
| 6 | 413 | 18/18 | 0.017 | 1.61 | 0.017 | UP* |
| 7 | 419 | 18/18 | 0.031 | 1.83 | 0.031 | UP# |
| 8 | 420 | 18/18 | 0.39 | 1.28 | 0.39 | UP# |
| 9 | 407 | 18/18 | 0.20 | 1.36 | 0.20 | UP# |

### APN2 expression in Cry toxin exposed larval midgut epithelium

*In situ* hybridisation based APN2 transcript expression analysis in toxin exposed insects (G6) was carried out. Presence of DIG labelled APN cRNA was confirmed in transverse sections (Fig 7). The study revealed localization of APN2 transcript primarily localized in differentiated epithelial cells present towards lumen side. Decreased fluorescent visualized for APN2 cRNA in toxin exposed (G6) larval midgut further confirmed declined APN2 mRNA expression (Fig 7).

**Figure 7.**
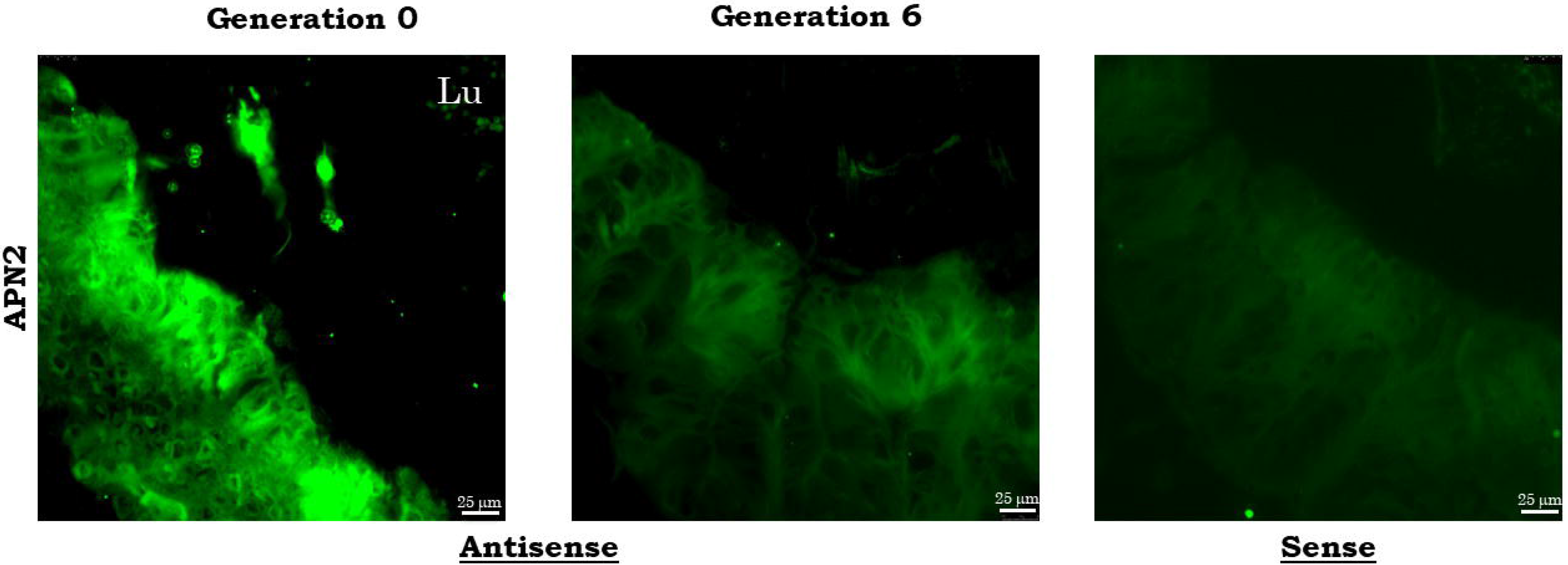
*In situ* APN2 mRNA expression analysis in toxin exposed larval midgut. Green fluorescence represents the presence APN2 transscript tagged with DIG labelled cRNA (→). Micrograph clearly shows that APN2 mRNA in G6 comparatively lower than G0.

## Discussion

Midgut is truly a multifaceted tissue which performs a variety functions including digestion, absorption, immune function, transmission of pathogens and interaction with various ingested molecules (Kliot and Ghanim 2016; Kim et al., 2016; Wu et al., 2016). Fairly large numbers of proteins present in the apical brush border of midgut epithelial cells are either reported as digestive proteases or transporters which facilitate the aforesaid functions. Among them proteins like cadherins (Nagamatsu et al., 1998; Walsh et al., 2018), GPI-anchored APNs (Knight et al., 1995; Rajagopal et al., 2003;de Bortoli and Jurat-Fuentes, 2018; Sun et al., 2019), ALPs (McNall and Adang, 2003; Jurat-Fuentes and Adang, 2004; Ren et al., 2018) and ABCC2 (Tanaka et al., 2013; Zhou et al., 2016; Nakaishi et al., 2018) present in brush border were shown to interact with Cry toxin(s) produced by several strains of *B. thuringiensis* and were characterized as Cry toxin functional receptor in midgut epithelium of insects.

Cry toxin(s) produced by DOR *Bt*-1 strain were shown to have high insecticidal activity towards the larval forms of *A. janata* as well as few other lepidopteran pests (Vimala Devi et al., 2005; 2006). Ligand binding studies in the present study revealed interaction of five to six midgut epithelial cell membrane proteins with the activated toxin(s) obtained from DOR *Bt*-1 as well as recombinant Cry1Ac (BGSC). However, prominent interaction was primarily seen with a ~110 kDa protein which was confirmed as APN by immunodetection using anti-APN antibody. MALDI-TOF/TOF analyses further confirmed that Cry interacting protein (~110 kDa) in *A. janata* BBMVs is aminopeptidase N. Colocalization studies substantiate the presence of APN and Cry toxin in close proximity in apical brush border membrane of midgut epithelium. The results obtained in present study corroborate well with earlier reports where brush border membrane region was shown to interact and accumulate toxins in intoxicated larvae (Bravo et al., 2007; Yi et al., 1996; Valaitis, 2011; Onofre et al., 2017). Reported molecular weights of APNs in different insects ranged between 100-150 kDa. Multiple isoforms have been reported in various lepidopterans, *B. mori* (Nakanishi et al., 2002)*, T. ni* (Wang et al., 2005), *O. nubilalis* (Crava et al., 2013) and *H. virescens* (Perera et al., 2015). The screening of midgut transcriptome data generated from our laboratory (SRR7212215) also revealed presence of five different isoforms of APN in larval midgut of *A. janata* (Dhania et al., 2019). They were cloned and confirmed by Sanger sequencing. *In silico* analysis of identified sequences revealed the presence of characteristic aminopeptidase activity motif “GAMENEG” and zinc-binding motif “HEXXHX_18_H”. The identified isoforms have been named, based on the homology found with reported sequences in NCBI database as APN1 (3054 bp, 1017aa, ~114 kDa, pI 4.73); APN2 (2835 bp, 944 aa, ~107 kDa, pI 4.81); APN4 (2853 bp, 950 aa, ~108 kDa; pI 4.81); APN6 (2868 bp, 955aa, ~109 kDa, pI 4.94) and APN9 (2280 bp; 759 aa, ~88 kDa, pI 5.43). The molecular weight of all the isoforms of APN identified in *A. janata* ranged between 100-120 kDa, except APN9 which was a low molecular weight protein around 88 kDa. A low molecular weight APN of 90 kDa was also reported in *B. mori* (Nakanishi et al., 2002). Further, the five isoforms cloned from larval midgut of *A. janata* showed a maximum of 37% sequence identity with each other. Detailed analyses confirmed that the identified isoforms were paralogs of each other and were expressed by different genes of single cluster. Phylogenetic review also suggested that most of the APNs were derived by gene duplication, from a recent common ancestor of lepidopteran and dipteran gene (Hughes, 2014). In addition, conserved domain analysis of the identified sequences revealed that they belonged to M1 aminopeptidase N family with ERAP1 [M1_APN_2 and ERAP1_C domain-containing protein], which were type2 integral membrane proteases (Kola et al., 2016). *In silico* analyses showed the presence of transmembrane peptide at C-terminal end of APN1, APN2 and APN4, in addition APN1 and APN4 showed signal peptide at N-terminal which was absent in APN2. On the other hand, APN6 showed signal peptide at N-terminal but not the transmembrane peptide at C-terminal. It was interesting to note that APN9 isoform cloned from larval midgut of *A. janata* lacked both C-terminal as well as N-terminal signal sequences.

In local fields, for the management of *A. janata*, sprays of the *Bt* formulations are applied with a gap of three days for the control of late 2nd or early 3rd instar larvae. The same strategy was followed in the laboratory to generate Cry toxin exposed insects to analyze the expression profile of APN isoforms. In a recent study, we have clearly demonstrated that even at sublethal dosage, Cry toxin induced extensive cell death in larval midgut, promoted mortality and caused defective metamorphosis (Chauhan et al., 2017). In the present study APN expression profiling was carried out in sublethal Cry toxin exposed larval populations (generation wise analysis). It was interesting to note that continuous exposure of toxin for 72 h, caused a gradual decline in all APN isoforms and each isoform was differentially regulated. On the other hand, the expression profile of APN isoforms from generation wise analyses of toxin exposed larval populations (for 6 continuous generations) revealed the declined expression of APN2 and inclined expression of APN4. These findings suggest that upon Cry toxin exposure, larval midgut has ability to modulate the expression of APNs. Tiewsiri and Wang, (2011) have already reported down-regulation of one of the isoform (APN1) in Cry resistant larvae of cabbage looper, *T. ni*. In present study, DIGE expression analysis carried out with BBMV proteins isolated from G0 and G6 larvae, revealed up-regulation of most of the protein spots which were detected using anti-APN antibody. Result showed that there was one protein spot which was slightly down-regulated; an analysis of this protein spot using MALDI-MS/MS analysis confirmed it as APN2. *In situ* hybridization studies confirmed lower expression of APN2 in toxin exposed sixth generation. Further for 6th generation of toxin exposed insects with respect to first generation and increase in LC50 value from 247.2 μg/ml to 298.47 μg/ml was also seen.

Along with above findings and the *in silico* analysis which revealed absence of signal peptide in APN2 and presence of transmembrane domain once again suggested that APN2 in *A. janata* is a membrane-bound aminopeptidase N. As Cry toxin interaction with receptor protein is known to facilitate the pore formation leading to damage and cell death (Bravo et al., 2004: Soberon et al., 2009). Reduced expression of APN2 seen in the present study might lower the interaction of midgut apical brush border with Cry toxin(s). Findings of the present study suggest that down-regulated APN2 which might act as a Cry toxin receptor in *A. janata* larvae and might be responsible for lowering the cellular damage and facilitated toxin tolerance.

## Conclusions

With the present finding of the study we propose APN2 act as one of the Cry toxin receptors which mediated Cry toxin based damage in larval forms of *A. janata*. The altered expression of APNs upon sublethal Cry toxin exposure during generation wise study might facilitate the toxin tolerance in larvae. Overall this study, suggests that *A. janata* larvae exploit a variety of strategies to avoid the deleterious effects of Cry intoxication.

## Acknowledgments

The work was carried out with financial support of DST-SERB grant (Grant No. SB/SO/AS-047/2013), and partially UGC-BSR Faculty Fellowship to ADG. DOR *Bt*-1 formation was a research gift from IIOR, Hyderabad. Financial support in form of research fellowship to VC by DBT and ND, VL, BB by UGC, India are acknowledged. The common central facilities and instruments generated with support of DST-FIST (Level-II), DBT-CREBB, UPE-II and DST-PURSE were used for the present study.

## Author contributions

V.C., N.D., V.L. and B.B. performed all experimental work. K.P. help in providing facility and analysis of 2D gel electrophoretic results. A.D. conceived, designed, analysed and drafted the manuscript. All authors have read and approved the final manuscript.

---

**Supplementary Figure 1. *In silico* analysis of APN isoforms.** (a) APNs from larval mid-gut show the presence of aminopeptidase active motif “GAMENEG” and zinc-binding motif “HEXXHX_18_H”. (b) The comparative study shows that APN isoforms of *A. janata* have only 37% identity. (c). Conserved domain analysis shows the presence of transmembrane peptide and signal sequence. APN1 and APN4 show the presence of transmembrane and signal peptides, APN2 shows only transmembrane peptide, APN6 shows N-terminal signal peptide while APN9 lacked both N and C-terminal peptides.

**Supplementary Figure 2. Identification of APN proteins.** Two dimension SDS-PAGE and immunoblotting with APN antibody showed the presence of multiple cross reacting isoforms of APNs (pI 4.8-5.4).

