## Supplementary Fig. 1 for "Midgut aminopeptidase N expression profile in Castor semilooper during sublethal Cry toxin exposure"

a. Multiple sequence alignment

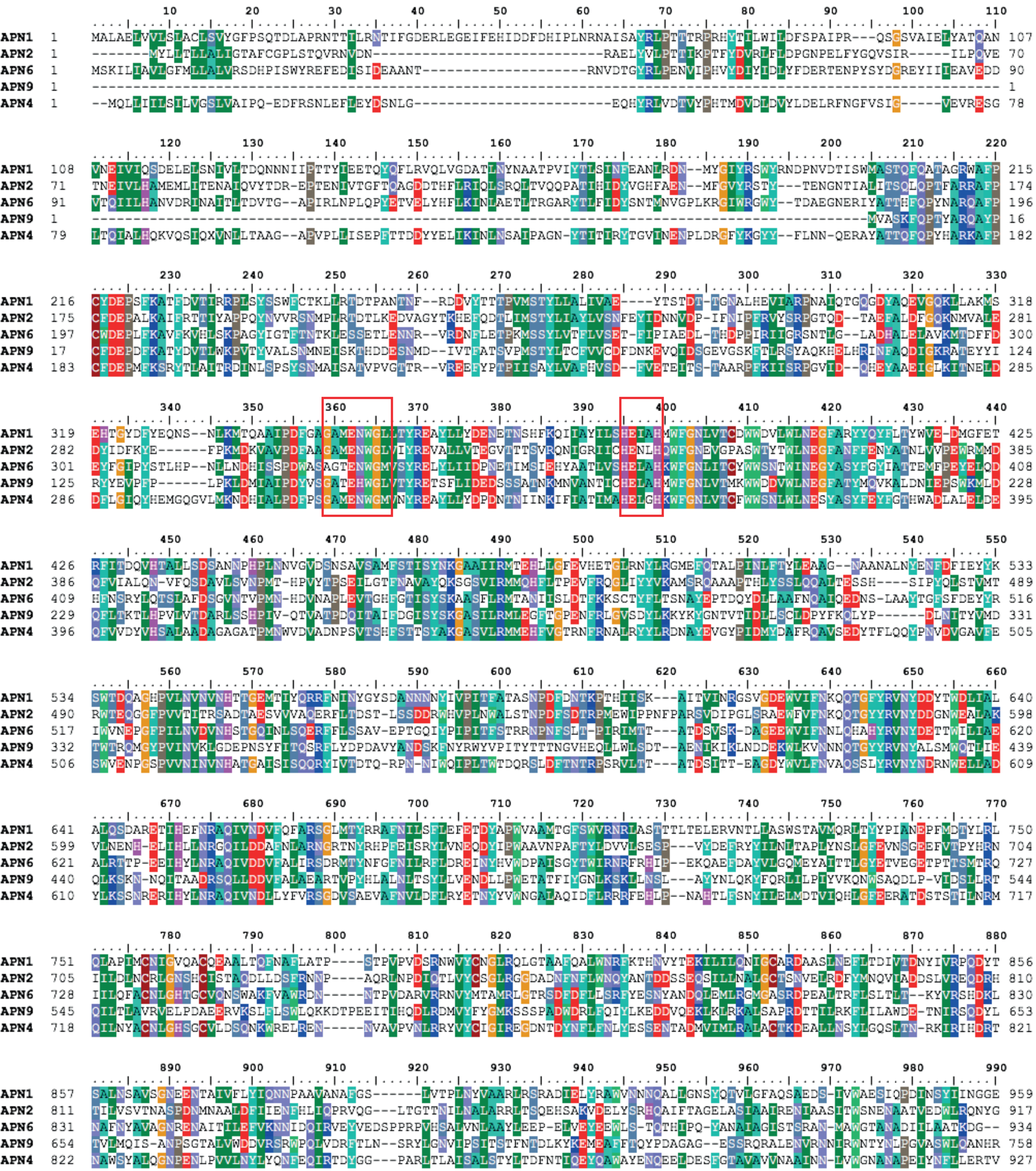

b. Percent Identity Matrix

| | | APN1 | APN2 | APN4 | APN6 | APN9 |
|---|---|---|---|---|---|---|
| 1 | APN1 | 100.00 | 29.15 | 31.22 | 30.87 | 27.50 |
| 2 | APN2 | 29.15 | 100.00 | 29.48 | 27.08 | 27.82 |
| 3 | APN4 | 31.22 | 29.48 | 100.00 | 37.43 | 26.09 |
| 4 | APN6 | 30.87 | 27.08 | 37.43 | 100.00 | 24.86 |
| 5 | APN9 | 27.50 | 27.82 | 26.09 | 24.86 | 100.00 |

c. Conserved domain analysis

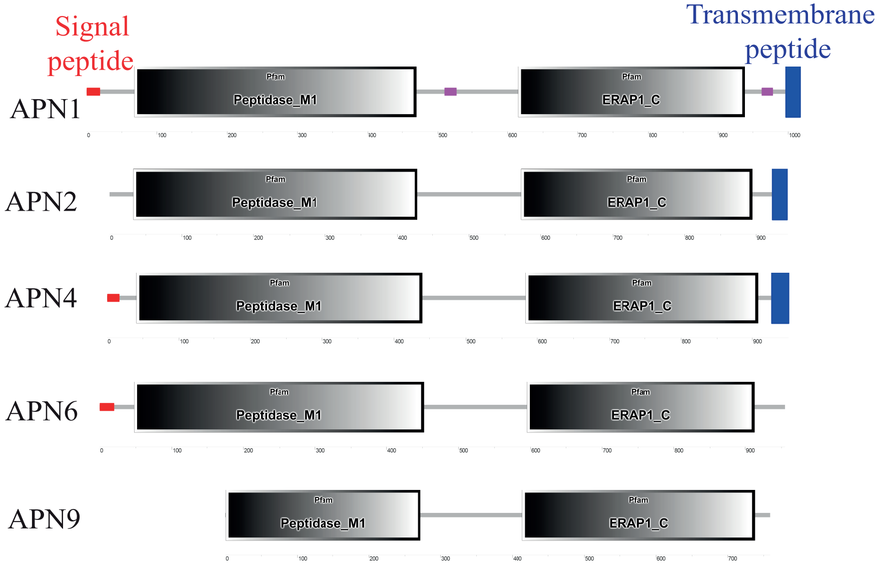
