## Supplementary figures and images for "Midgut aminopeptidase N expression profile in Castor semilooper during sublethal Cry toxin exposure"

### Supplementary Fig. 2

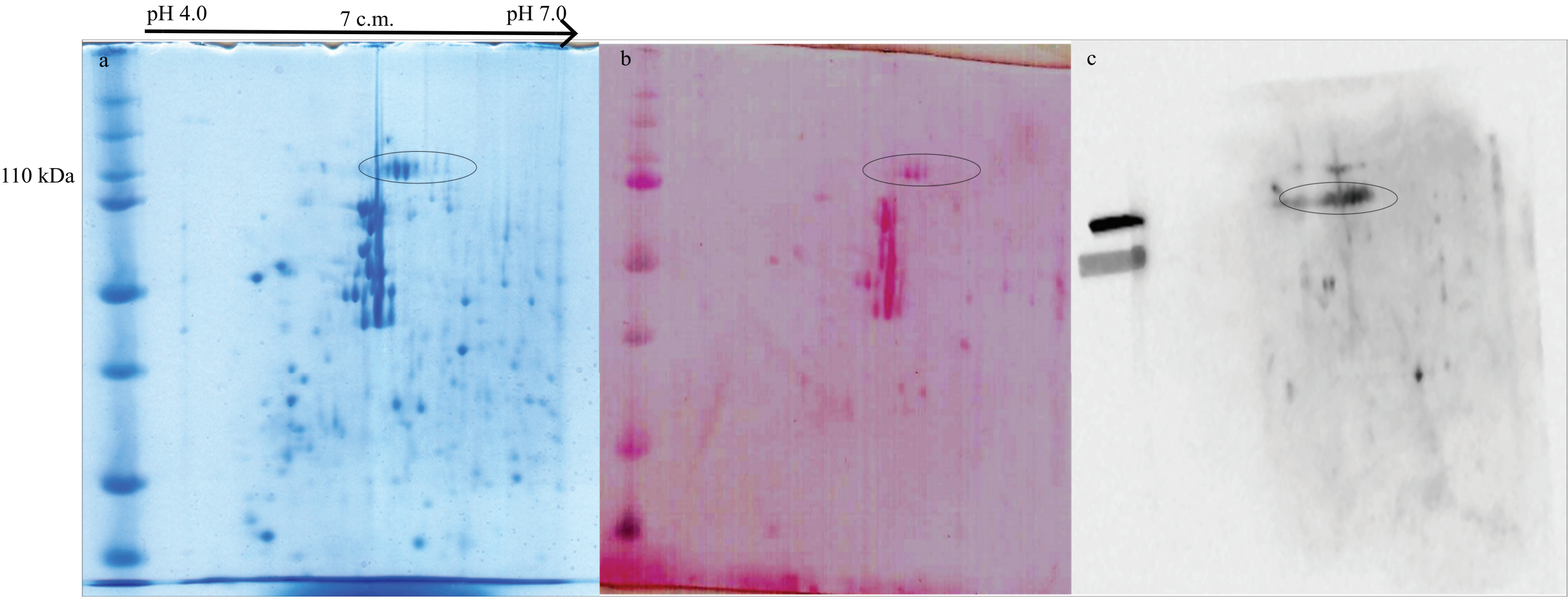
